# Effect of species, size, and chimerism on the susceptibility of Caribbean brain coral recruits to stony coral tissue loss disease (SCTLD)

**DOI:** 10.1101/2021.11.26.469756

**Authors:** Olivia M. Williamson, Caroline E. Dennison, Keri L. O’Neil, Andrew C. Baker

**Affiliations:** Department of Marine Biology and Ecology, Rosenstiel School of Marine and Atmospheric Science, University of Miami, 4600 Rickenbacker Cswy., Miami, FL, 33149, USA; Center for Conservation, The Florida Aquarium, Apollo Beach, FL, USA

**Author notes:** **Correspondence:** Olivia Williamson.

**Keywords:** coral, recruits, stony coral tissue loss disease, *Diploria labyrinthiformis*, *Colpophyllia natans*, chimera, resilience

## Abstract

Stony coral tissue loss disease (SCTLD) has devastated coral populations along Florida’s Coral Reef and beyond. Although widespread infection and mortality of adult colonies have been documented, no studies have yet investigated the susceptibility of recruits to this disease. Here, we exposed eight-month-old *Diploria labyrinthiformis* recruits and four-month-old *Colpophyllia natans* recruits to two sequential doses of SCTLD in the laboratory to track infection and assess potential resilience. Both species began to develop lesions as early as 48 h after exposure began. During the first dose, 59.0% of *C. natans* recruits lost all tissue (died) within two to eight days of developing lesions, whereas *D. labyrinthiformis* recruits experienced significantly slower rates of tissue loss and minimal eventual mortality. In *C. natans*, larger recruits and those fused into groups of multiple genets (chimeras) exhibited the highest survivorship. In contrast, smaller and/or single (ungrouped) recruits had the lowest survivorship (9.9 - 26.5%). After 20 days, a second SCTLD dose was delivered to further test resistance in remaining recruits, and all recruits of both species succumbed within 6 days. Although no recruits showed absolute resistance to SCTLD following repeated exposures, our results provide evidence that interactions between species, size, and chimerism can impact relative resistance. This study represents the first report of SCTLD in Caribbean coral recruits and carries implications for natural species recovery and reef restoration efforts. Additional research on the susceptibility of coral juveniles to SCTLD is urgently needed, to include different species, locations, parents, and algal symbionts, with the goal of assessing relative susceptibility and identifying potential sources of resilience for this critical life history stage.

## 1 Introduction

Stony coral tissue loss disease (SCTLD) has devastated coral reefs in Florida since its first outbreak near Miami in 2014. This disease has been found to infect at least 22 species of scleractinian corals, making it one of the most lethal coral disease outbreaks ever recorded (Precht et al., 2016). It has spread rapidly throughout Florida’s Coral Reef and beyond, resulting in extremely high rates of infection and mortality at affected sites (Precht et al., 2016; Walton et al., 2018; FKNMS, 2018; Alvarez-Filip et al., 2019; Sharp et al., 2020; Dahlgren et al., 2021; Brandt et al., 2021). Localized outbreaks have precipitated significant declines in live coral tissue area (>60%) over short periods of time (Walton et al., 2018; Sharp et al., 2020; Heres et al., 2021; Brandt et al., 2021), with some species being more severely impacted than others (Gintert et al., 2019; Costa et al., 2021; Neely et al., 2021a; Spadafore et al., 2021). In surveys in Miami-Dade county in 2015 and 2016, Precht et al. (2016) documented 83% infection in *Diploria labyrinthiformis* colonies, and estimated that these populations were reduced to <25% of their original numbers. Such high rates of infection and mortality may jeopardize the long-term persistence of susceptible species (Precht et al., 2016; Neely et al., 2021a). Furthermore, these dramatic losses have altered the composition of coral communities in affected areas, threatening biodiversity and ecosystem function (Alvarez-Filip et al., 2013; Walton et al., 2018; Gilliam et al., 2019; Estrada-Saldívar et al., 2020; Sharp et al., 2020; Heres et al., 2021).

The impacts of SCTLD have been well documented in many locations experiencing outbreaks, but details on the cause(s) and dynamics of this disease remain unknown. Although the causative pathogen(s) for SCTLD remain unidentified, enrichment of disease-associated bacteria in lesions, along with the efficacy of antibiotics, suggests that bacteria play a role in disease progression (Meyer et al., 2019; Neely et al., 2020; Rosales et al., 2020; Neely et al., 2021b; Clark et al., 2021). Transmission has been shown to occur through direct contact as well as through the water column in neutrally buoyant particles (Aeby et al., 2019; Dobbelaere et al., 2020; Eaton et al., 2021). Williams et al. (2021) found that in the Florida Keys, SCTLD disproportionately affected large corals and areas with high species diversity and colony density. In an experimental exposure Dennison et al. (2021) found that corals manipulated to contain algal symbionts in the genus *Durusdinium* were less susceptible to SCTLD compared to their counterparts hosting *Breviolum* and *Cladocopium*, and Rubin et al. (2021) reported that inshore brain corals hosting *Durusdinium* largely escaped infection compared to offshore corals hosting *Breviolum*. Beyond these reports, however, little is known about the factors contributing to infection and/or resilience in coral colonies.

Reef surveys tracking disease prevalence in SCTLD-susceptible species have only documented the health of colonies ≥4 cm in diameter, excluding recruits (Gilliam et al., 2019; Kramer et al., 2019). Consequently, we have no knowledge of the extent to which Caribbean coral juveniles may have been impacted by SCTLD, nor the degree of risk they face from future outbreaks. If populations of susceptible species are to recover, it will be important to understand whether new generations of corals (i.e., sexually-produced juveniles) that recruit to the reef are susceptible to disease. In addition, as scientists and managers test active reef restoration techniques to restore depleted populations, they will need to consider the danger that SCTLD poses to outplanted corals (both asexually-produced fragments and sexually-produced recruits).

Even in the absence of environmental stressors such as disease, the early life stages of many corals are characterized by high mortality and population “bottlenecks” that severely reduce the number of living recruits over time (Vermeij & Sandin, 2008). A recruit’s tiny size leaves it vulnerable to many organisms on the reef capable of preying on, grazing over, outcompeting, or simply smothering it (Doropoulos et al., 2012; 2016). As such, recruits benefit from growing as quickly as possible, becoming large enough to be less susceptible to predation and competition (Doropoulos et al 2012, 2016). In addition, larvae often settle in groups of multiple individuals, and these aggregations may have implications for survival during early ontogeny (Duerden, 1902; Frank et al., 1997; Raymundo & Maypa, 2004; Puill-Stephan et al., 2011, 2012). In particular, the fusion of multiple primary polyps into a chimeric colony may allow vulnerable coral recruits to rapidly surpass size-escape thresholds (Raymundo & Maypa, 2004; Amar et al., 2008; Christiansen et al., 2008; Vermeij & Sandin, 2008; Puill-Stephan et al., 2011; Doropoulos et al., 2016; Sampayo et al., 2020). Moreover, the inherent genetic diversity of chimeric colonies is also thought to offer advantages for coral resilience (Rinkevich & Yankelevich, 2004; Rinkevich et al., 2016; Rinkevich, 2019; Huffmyer et al., 2021). With high mortality already a hallmark of early ontogeny in corals, it is likely that SCTLD outbreaks may exacerbate the bottlenecks that impact populations of juvenile corals.

To date, there are no reports on whether Caribbean coral recruits can become infected with SCTLD or how their pathology may compare to what has been observed in adult colonies. Here, we exposed four-month-old *Colpophyllia natans* and eight-month-old *Diploria labyrinthiformis* recruits to SCTLD by rearing them for four weeks in semi-recirculating laboratory systems shared with infected adult colonies of four different coral species and monitoring patterns of survival and disease progression. We tracked the fate of each recruit throughout exposure in relation to its size and recruit grouping status (chimera vs. non-chimera), and characterized susceptibility by lesion prevalence, lesion progression rates, and mortality.

## 2 Methods

### 2.1 Gamete collection

This study utilized two cohorts of coral recruits collected as gametes from parent colonies in *ex-situ* induced spawning facilities at the Florida Aquarium’s Center for Conservation. In both cases, parent colonies had been collected by the Florida Fish and Wildlife Conservation Commission as part of the Coral Rescue Project (under FKNMS Superintendent permit FKNMS-2017-100), after which they were loaned to The Florida Aquarium to serve as a living genetic archive. On May 20, 2020, eight *Diploria labyrinthiformis* colonies from the Marquesas and Key West spawned, and their gametes were pooled to make a cohort of offspring. On September 9, 2020, three *Colpophyllia natans* colonies spawned, and their gametes were pooled to make a cohort of offspring.

Two days post-fertilization, developing larvae were transported to an indoor laboratory at the University of Miami’s Rosenstiel School of Marine and Atmospheric Science and reared in UV-sterilized, one-micron filtered seawater (FSW), with temperature maintained at the ambient level of incoming water from Biscayne Bay (~26-30°C in spring-summer). Each batch of larvae was settled on unconditioned 2.5-cm ceramic plugs (Boston AquaFarms, USA). After five days, settlers were counted under a microscope.

### 2.2 Recruit development

After settlement, recruits were provisioned with Symbiodiniaceae. Fragments of *Orbicella faveolata* hosting *Durusdinium trenchii* were placed in aquaria with *D. labyrinthiformis* recruits to serve as symbiont sources (Williamson et al., 2021). The *C. natans* recruits were inoculated with approximately one million cells per liter of cultured *Breviolum* and *D. trenchii* twice per week, until all recruits were visibly infected (approximately three weeks). For the next several months, no further Symbiodiniaceae sources were provided, and recruits were fed twice per week with Reef Roids (PolypLab). Temperature tracked local seasonal fluctuations of incoming seawater in Bear Cut, FL, eventually reaching 22°C in January 2021.

At this point, a subset of recruits from each cohort were sampled to characterize their Symbiodiniaceae communities. Briefly, small tissue biopsies (<0.25 cm^2^) were taken from each recruit using a razor blade, genomic DNA was extracted following modified organic extraction methods (Rowan and Powers, 1991; Baker & Cunning, 2016), and quantitative PCR (qPCR) assays were used to identify Symbiodiniaceae to genus level following reactions described in Cunning and Baker (2013) using a QuantStudio 3 Real-Time PCR Instrument (Applied Biosystems, USA). All *D. labyrinthiformis* recruits and all but two *C. natans* recruits hosted primarily (>90%) *D. trenchii*. One *C. natans* recruit hosted 10.4% *Breviolum* and 89.6% *D. trenchii*, and one hosted *Breviolum* only.

After sampling, recruits from each cohort were counted, photographed, and measured under a microscope. Because many recruits had settled in clusters, we recorded whether each recruit was part of a group when assessing its susceptibility to SCTLD. If a recruit grew to contact another to form a multi-genotype entity, it was deemed a member of a “chimera” (Fig. 1 A, D). In all instances of tissue contact, fusion occurred to form stable chimeras; no evidence of nonfusion or incompatible fusion were observed (Frank et al 1997; Nozawa & Loya 2005). If a recruit was isolated from other recruits with no physical contact, it was deemed “single”. ImageJ was used to measure maximum diameter and recruit skeletal area in mm^2^. Each recruit was also categorized by size, based on its maximum diameter (*C. natans*: “small” = <1 mm, “medium” = 1-5 mm, “large” = >5 mm [Fig. 1 A-C]; *D. labyrinthiformis*: “small” = <5 mm, “medium” = 5-10 mm, “large” = >10 mm [Fig. 1 D-F]). These size and grouping categories were then tracked throughout the subsequent experiment.

**Figure 1:**
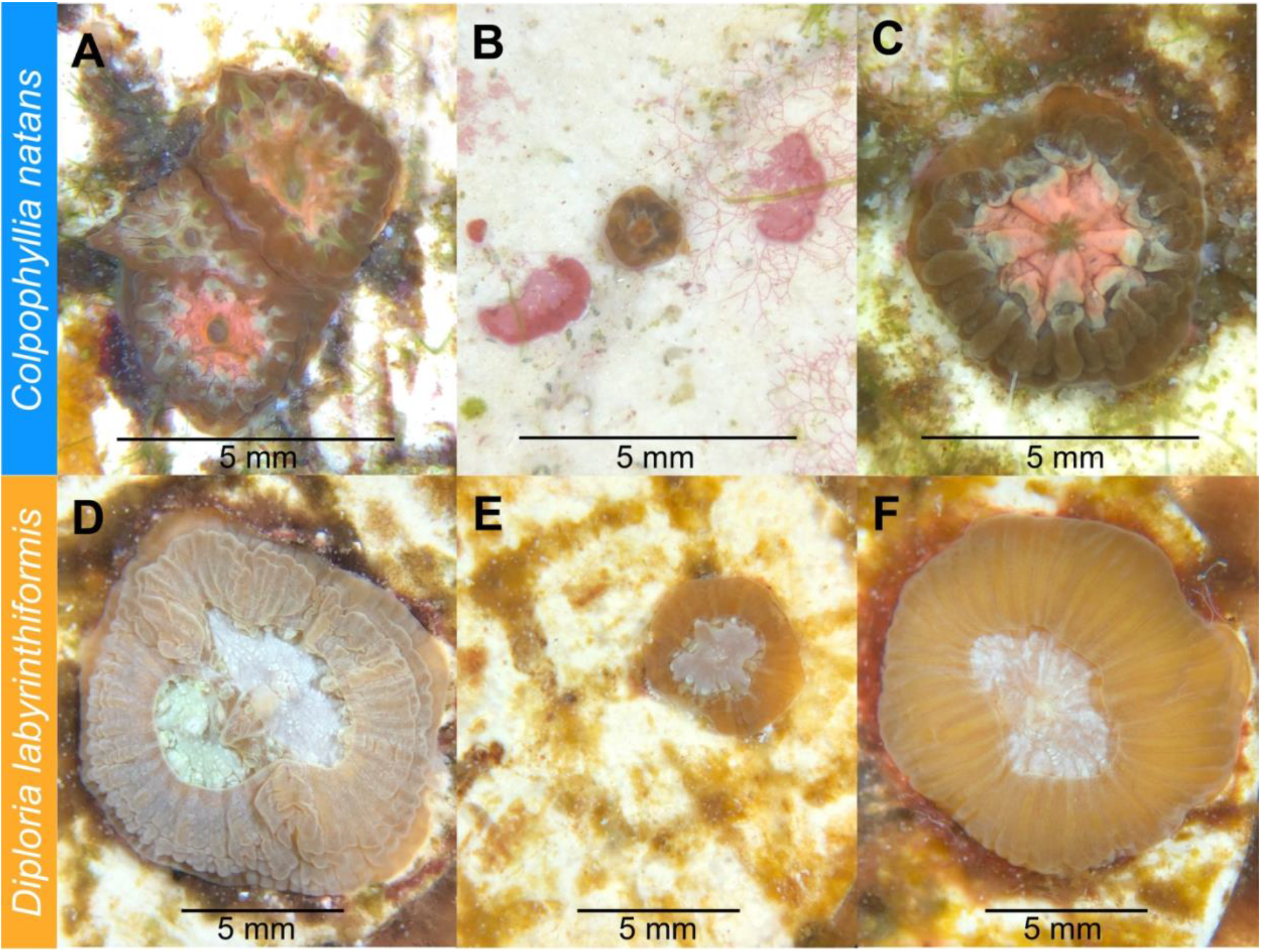
Four-month-old *Colpophyllia natans* **(A-C)** and eight-month-old *Diploria labyrinthiformis* (**D-F**) recruits prior to SCTLD exposure. **(A)** *C. natans* “chimera” (multi-genotype entity), made up of individual recruits ranked as “medium” in size. **(B)** *C. natans* “solitary” recruit (single-genotype entity), ranked as “small” in size. **(C)** *C. natans* solitary recruit, ranked as “large” in size. **(D)** *D. labyrinthiformis* chimera, made up of two individual recruits ranked as “medium” and “large” in size. **(E)** Small, solitary *D. labyrinthiformis* solitary recruit. **(F)** Large, solitary *D. labyrinthiformis* recruit.

### 2.3 SCTLD exposure

The ceramic plugs on which each species had settled were then distributed randomly into four new 2.5-gallon aquaria, so that each contained approximately 100 *C. natans* recruits and ten to twelve *D. labyrinthiformis* recruits, with all size and grouping categories represented (Fig. 2). As before, all aquaria were immersed in a larger 50-gallon seawater tank to maintain temperatures at ambient temperature of incoming water from Biscayne Bay (~22°C in January) to mimic what recruits would experience on the reef. Light (50–70 μmol quanta m−2 s−1, measured by an Apogee Underwater Quantum PAR Meter MQ-210) was maintained on a 12h:12h light:dark cycle using AI Hydra lights (AI Inc., USA). Irradiance and temperature were recorded with a HOBO Pendant® data logger (Onset Computer Corporation MX2202).

**Figure 2:**
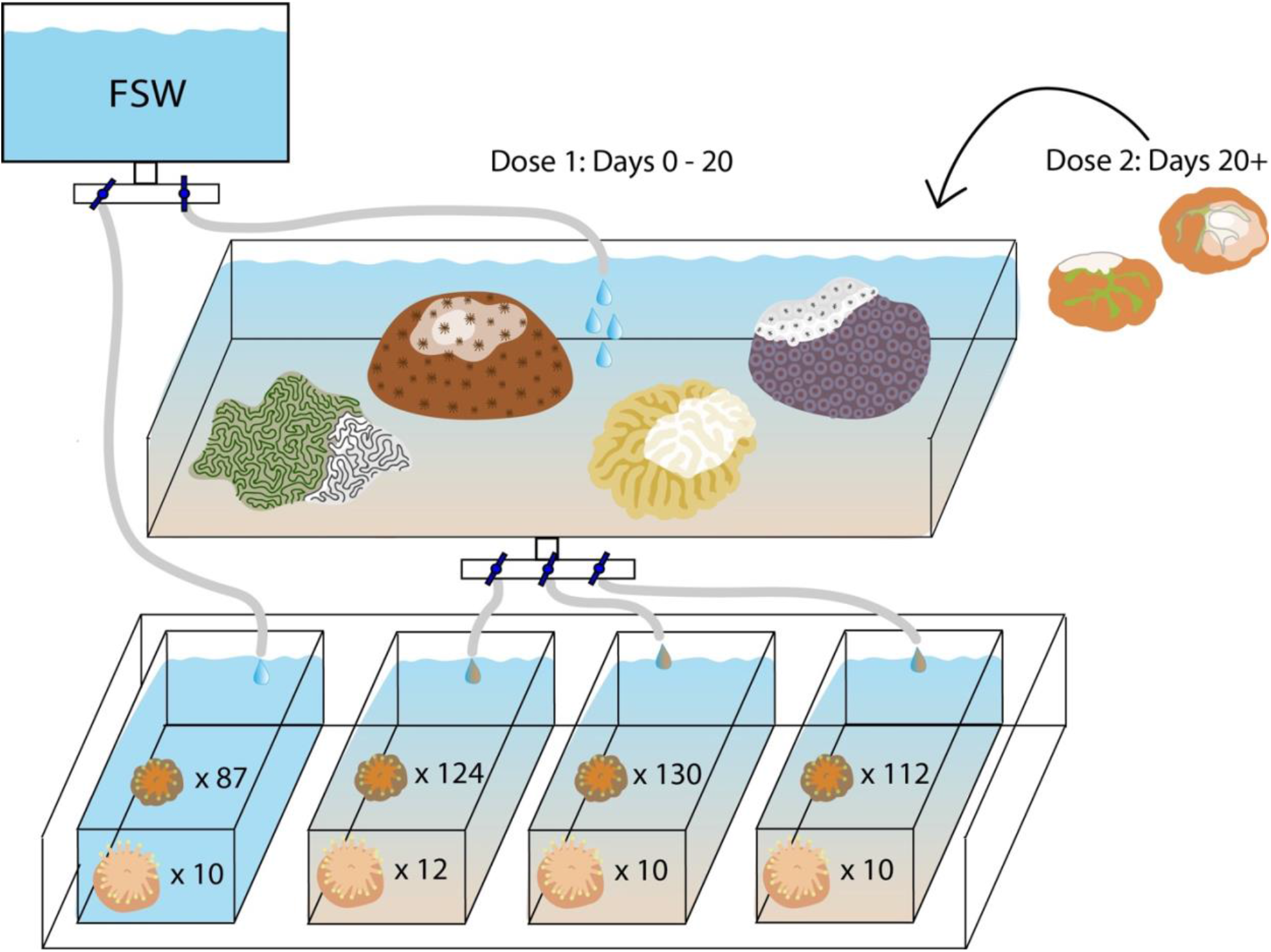
Experimental setup exposing eight-month-old *Diploria labyrinthiformis* recruits (n=42) and four-month-old *Colpophyllia natans* recruits (n=453) to SCTLD.

Three aquaria were supplied with seawater from another tank containing colonies of multiple species with active SCTLD lesions (approximately ~30 cm^2^ of tissue in total infected with SCTLD), which were sloughing off tissue and mucus that were presumed to contain the disease causative agent (Dobbelaere et al., 2020). This treatment was designed to replicate the waterborne nature of SCTLD transmission (Precht et al., 2016; Aeby et al., 2019; Muller et al., 2020), the most probable way by which recruits would become infected. Species present in this “disease bath” included *Pseudodiploria clivosa*, *Orbicella faveolata*, *Meandrina meandrites*, and *Siderastrea siderea*, each of which presented with varying degrees of SCTLD severity (FKNMS, 2018; Landsberg et al., 2020; Meiling et al., 2021). Water was dripped from the disease bath at 5L per hour into three of the aquaria, while the fourth (control) aquarium was supplied with FSW at the same rate (Fig. 2). One 4W submersible pump (VicTsing CAAA3-HG16) was placed into each aquarium to circulate water evenly.

While recruits were maintained in their respective treatments, algae were manually removed from plugs once per week to prevent overgrowth and/or competition. Recruits and diseased colonies were rotated within their respective tanks daily to avoid proximity bias. During the exposure, a Zeiss Stemi 305 LAB microscope (ZEISS International, Jena, Germany) was used to count the number of apparently healthy, infected, and dead recruits in each aquarium every other day. Pictures of each recruit and plug were taken under the microscope to document condition and lesion progression.

After 20 days, several small colonies (~5 cm in diameter) of *C. natans* with active lesions were added to the disease bath to deliver a second dose of SCTLD. This species has been reported to exhibit high rates of tissue loss (~10% per day, Meiling et al 2021), and colonies showed areas of bright white skeleton, indicating rapid disease progression. During this second exposure (hereafter referred to as “Dose 2”), recruits in each aquarium continued to be counted and photographed every other day.

Photographs from each time point were analyzed in ImageJ to measure the area of living tissue on affected recruits in mm^2^, which was standardized by initial recruit skeletal area measurements to calculate percent of living tissue remaining. Rates of tissue loss (% per day) were calculated by dividing the percent of tissue a recruit had lost at each time point by the number of days since it was last observed to be apparently healthy. R was used to run logistic regression models using the glm (generalized linear model) function comparing recruit survival and disease progression by species, size, and grouping (single vs. chimera). Models were fitted with logit links to account for binomial (survivorship) and quasibinomial (percent living tissue) distributions in the data.

## 3 Results

Overall, SCTLD gross morphology resembled descriptions provided by FKNMS (2018) and Landsberg et al. (2020). Starting 48 h after exposure began, some recruits of both species began to exhibit lesions. These typically began basally, sloughing tissue to expose denuded skeleton as they advanced across the recruit (Fig. 3). In *C. natans*, lesions were primarily focal, advancing from one area of tissue loss in a single front (Fig. 3). In some *D. labyrinthiformis* recruits, tissue loss presented multifocally, progressing from more than one initial lesion (Fig. 3).

**Figure 3:**
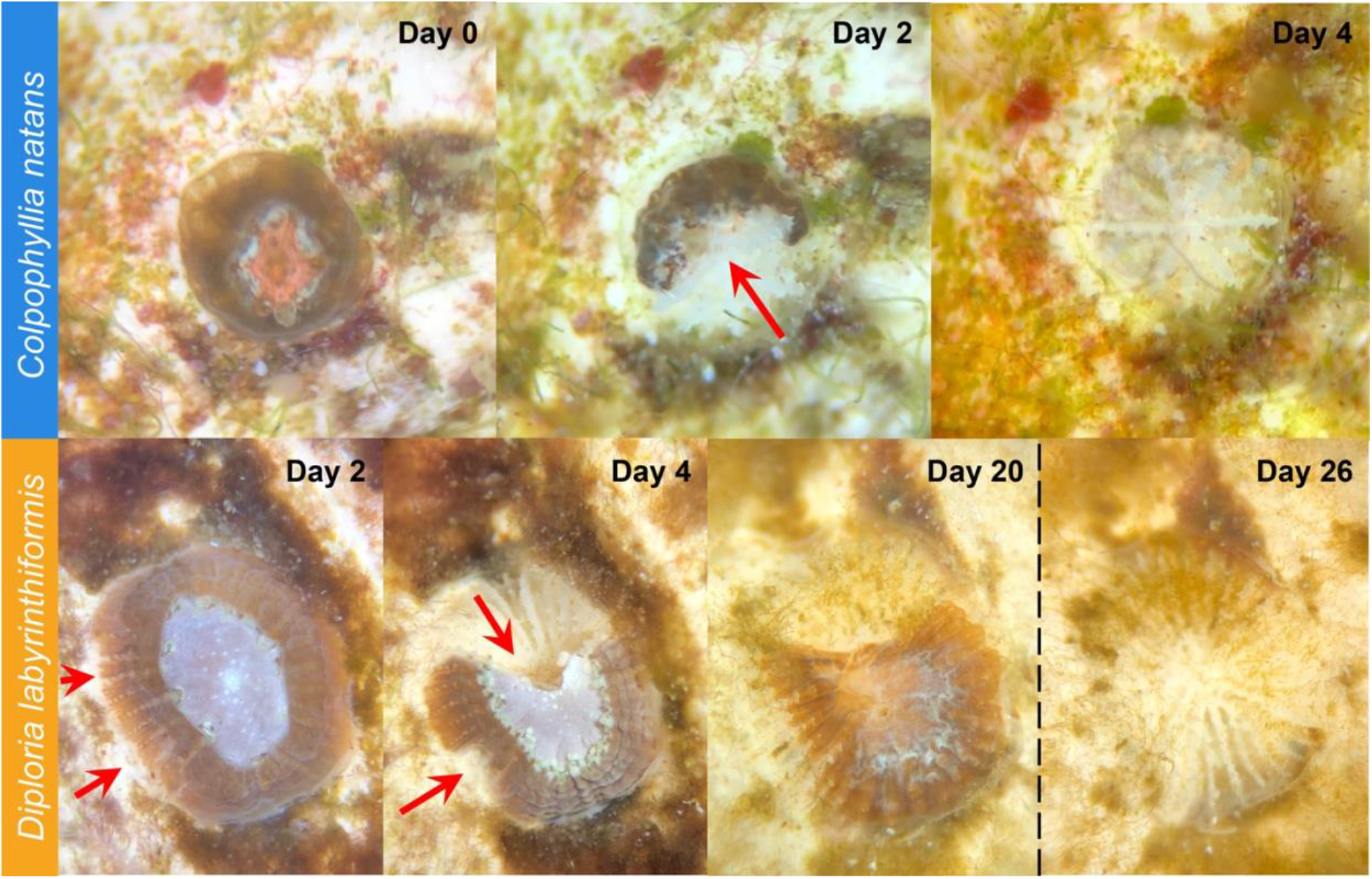
SCTLD progression in brain coral recruits. **Top:** *C. natans* recruit showing progression from apparently healthy to 100% tissue loss within four days (~25% tissue loss per day). Red arrow indicates active disease margins. **Bottom:** *D. labyrinthiformis* recruit showing progression from apparently healthy to ~50% tissue loss over four days, then halting lesion progression with no further major tissue loss until Dose 2 (dashed vertical line). Red arrows indicate active disease margins.

During the first 20 days of exposure (Dose 1), 64.8% of *C. natans* recruits in the disease treatments developed lesions and 59.0% died (Fig. 4). All *C. natans* recruits that became infected lost 100% of their tissue (died) within two to eight days after initial lesions (mean = 4.2 days, Fig. 5). Averaged across Dose 1, infected *C. natans* recruits lost 26.3 ± 13.6% of initial tissue per day (Fig. 5, 6). Five *C. natans* recruits in the control treatment (5.8%) died during the experiment without exhibiting signs of tissue loss, likely from algal overgrowth.

**Figure 4:**
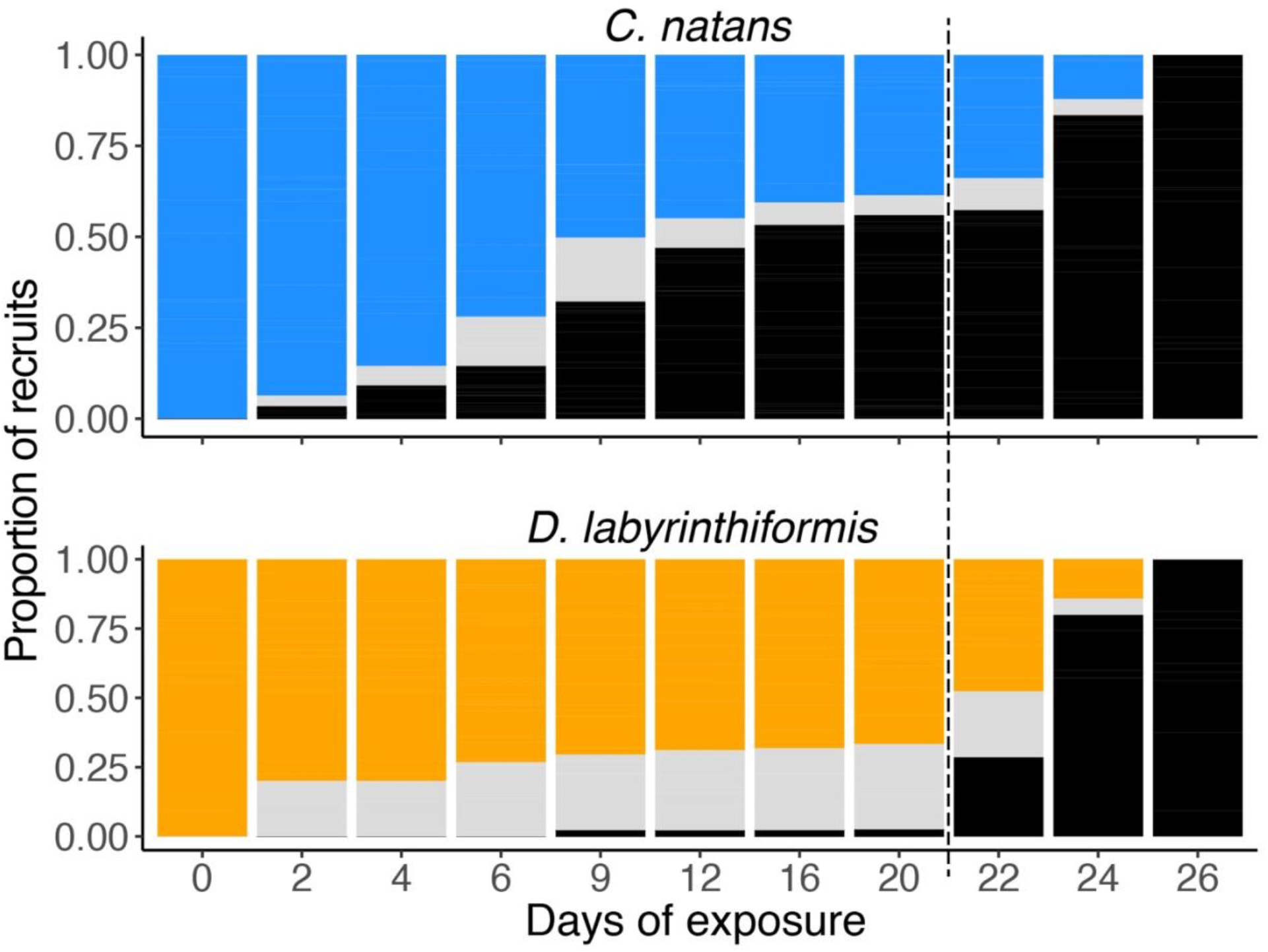
Condition of *C. natans* and *D. labyrinthiformis* recruits throughout SCTLD exposure. Blue (*C. natans*) and orange (*D. labyrinthiformis*) represent apparently healthy recruits, gray represents recruits exhibiting active disease lesions, and black represents recruits that have died. Dashed line indicates start of Dose 2.

**Figure 5:**
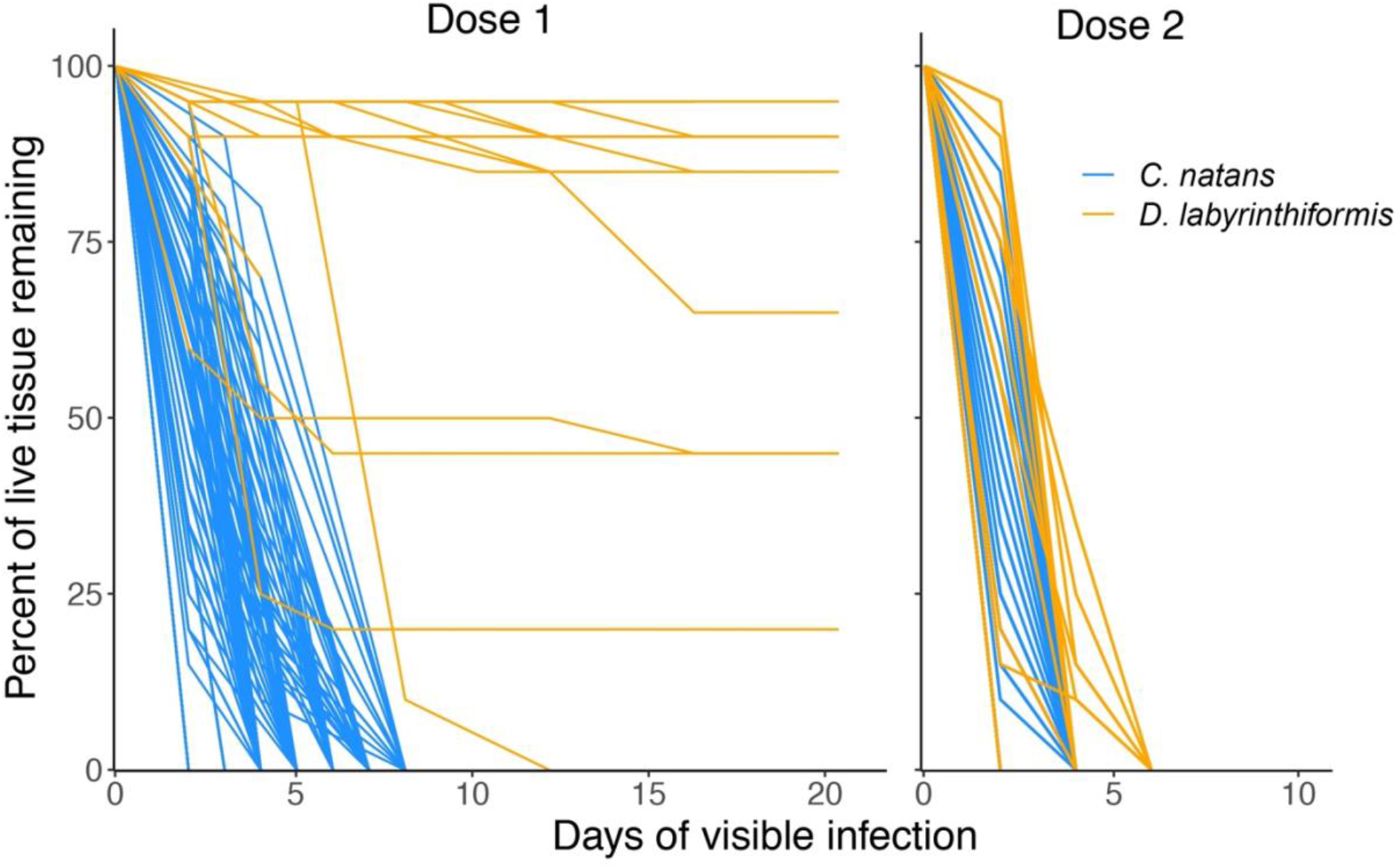
Loss of tissue over time as a result of SCTLD exposure in brain coral recruits. Lines represent individual recruits after developing lesions. Day 0 represents the last day that a given recruit was observed to be apparently healthy and have 100% of its tissue.

**Figure 6:**
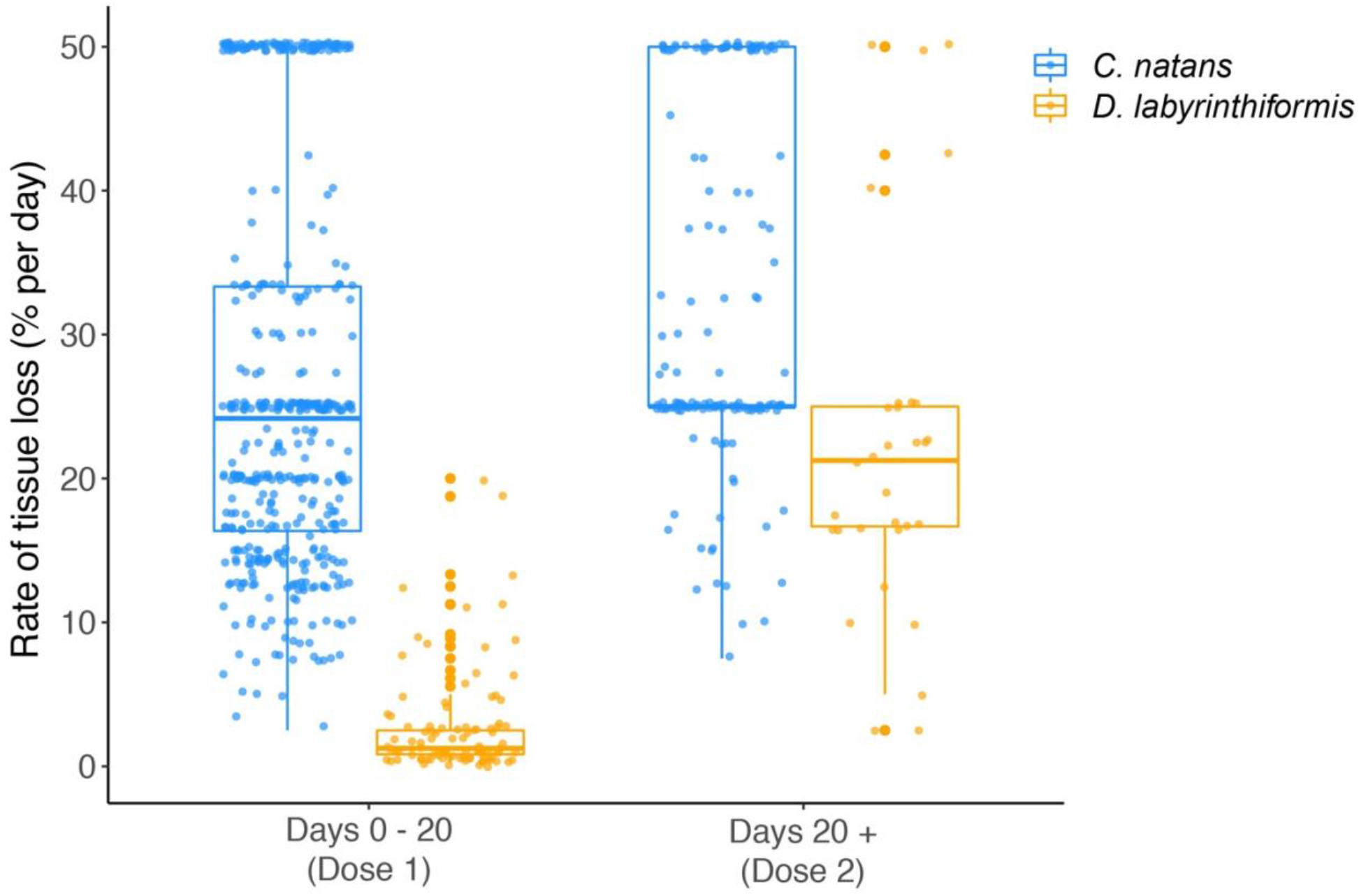
Rates of tissue loss as a result of SCTLD exposure in brain coral recruits. Points show measurements of tissue loss for individual recruits, and they were jittered to reduce overlap.

In contrast, 40.6% of disease-exposed *D. labyrinthiformis* recruits exhibited active disease lesions during Dose 1 (Fig. 4B), and only one recruit (3.1%) died (Fig. 4B). *D. labyrinthiformis* lost tissue at an average rate of 2.9 ± 3.7% per day, significantly slower than *C. natans* (p < 0.001, Fig. 5, 6). Most *D. labyrinthiformis* recruits experienced chronic necrosis (described by Landsberg et al. [2020] as “almost unnoticeable areas of recent necrosis apposed to skeleton overgrown with turf algae”) for the first three weeks of SCTLD exposure (Fig. 5). In several recruits, however, lesions progressed rapidly during the first four days of exposure, resulting in 45%-80% tissue loss, after which they slowed or stalled entirely (Fig. 3, 4B). None *D. labyrinthiformis* recruits in the control treatment exhibited tissue loss or died during the experiment.

In *C. natans*, Dose 1 revealed significant, interacting effects of recruit size and grouping on SCTLD susceptibility. Survivorship in *C. natans* recruits exposed to SCTLD increased significantly with polyp area (p < 0.001, Fig. 7A), and grouped individuals showed a significantly steeper increase in survivorship with size than single recruits (p < 0.001, Fig. 7A). For instance, a recruit that was 1 mm^2^ in area and grouped with others was ~30% more likely to survive than a single one of the same size, while 2 mm^2^ grouped recruits were ~50% more likely to survive than their single counterparts (Fig. 7A). In addition, we found a positive correlation between the number of *C. natans* genets (polyps) grouped together and the probability of survival during disease exposure (p < 0.001, Fig. 7B). Finally, when treated as a collective unit rather than individual recruits grouped together, small and medium-sized chimeras experienced significantly higher survivorship than single recruits of similar size (p = 0.0471 and p = 0.0076; Fig. 8). Overall, small, solitary recruits exhibited the lowest survivorship (9.9 ±3.2%), while large recruits in chimeric groups had the highest survivorship (100%, Fig. 8). In contrast, there were no significant differences in *D. labyrinthiformis* survivorship by size and grouping.

**Figure 7:**
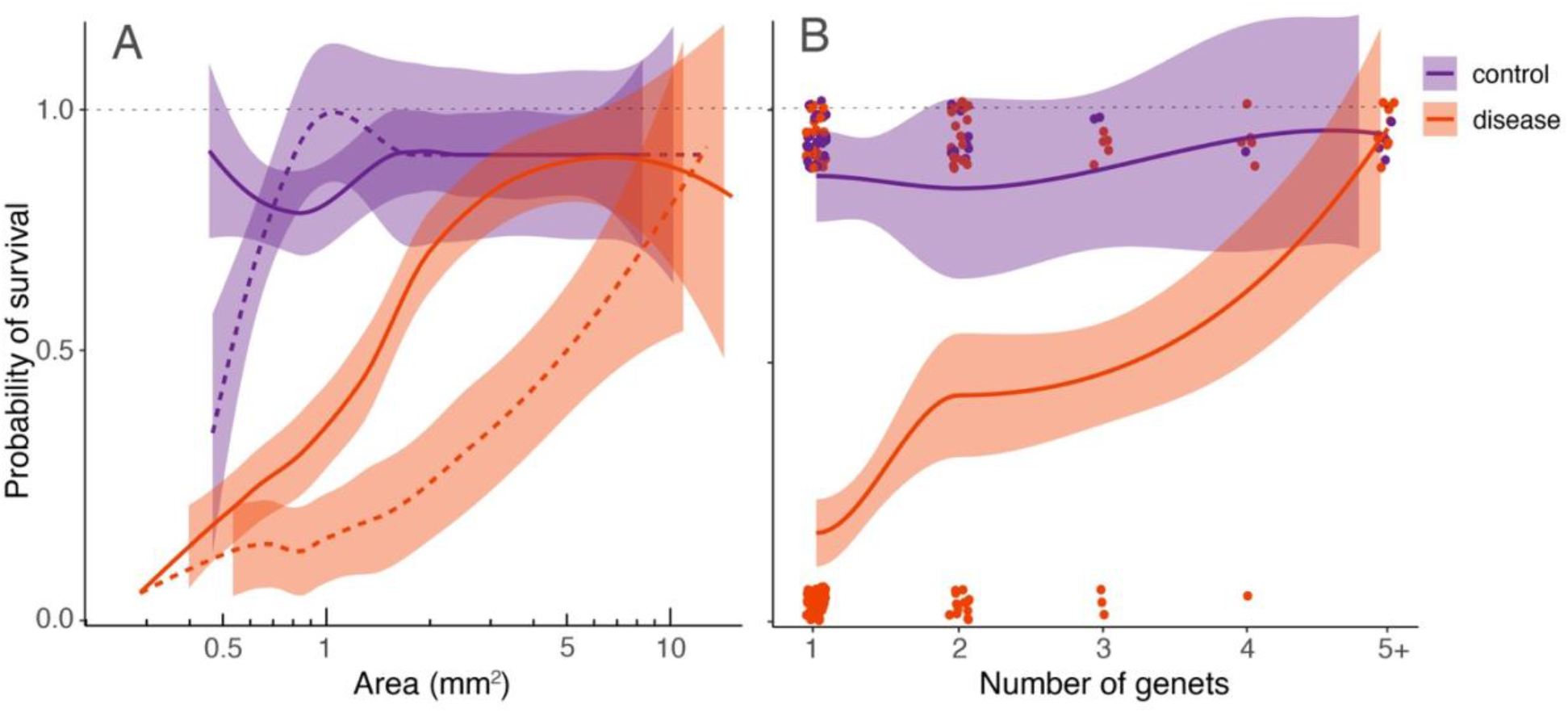
Relative susceptibility of four-month-old *C. natans* recruits to SCTLD during a 20-day experimental exposure (Dose 1). Recruits in the control treatment were not exposed to SCTLD, and thus represent the “background mortality” level expected for these juvenile corals. **A:** Mortality is negatively correlated with polyp area. Areal data were log(10) transformed to fit expectations of normalization. Dashed lines represent single recruits, while solid lines represent members of chimeric groups. **B:** Mortality is negatively correlated with number of genets/polyps grouped together. Points were jittered to prevent overlap. For both **A** and **B**, GLMs corrected for binomial survival data (method = “loess” [locally weighted scatterplot smoothing]) were used to fit the data. Shaded regions show 95% confidence intervals. Dotted line at y=1 to remind the reader of upper bounds (1).

**Figure 8:**
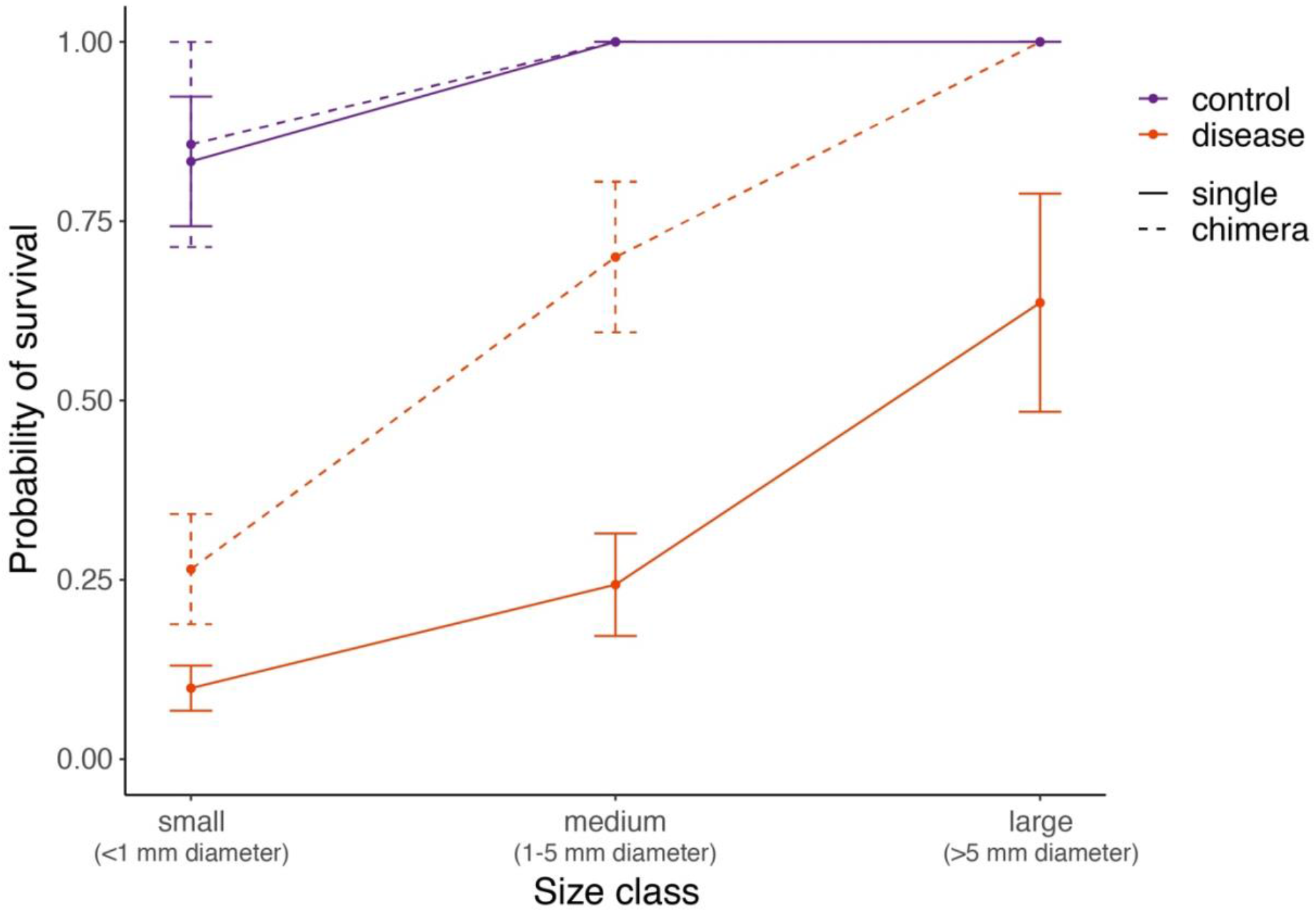
Relative SCTLD susceptibility of four-month-old *C. natans* varies by interactions of recruit size and grouping (single vs. chimera). Recruits in the control treatment were not exposed to SCTLD, and thus represent the “background mortality” level expected for these juvenile corals. A GLM with correction for binomial survivorship data (method = “loess” [locally weighted scatterplot smoothing]) was used to fit the data. Error bars indicate standard error.

Following the addition of new active SCTLD from several small *C. natans* colonies (Dose 2), all recruits of both species developed lesions and died within 6 days. This represented a significant increase in the rate of tissue loss in both species, from 2.9 ± 3.7% during Dose 1 to 22.1% ±12.7% during Dose 2 in *D. labyrinthiformis* (p < 0.001; Fig. 6) and from 26.3 ± 13.6% during Dose 1 to 33.0% ±12.8% during Dose 2 in *C. natans* (p < 0.001; Fig. 6).

## 4 Discussion

This study provides the first description of SCTLD pathology in Caribbean coral recruits. To date, recruits have not been reported in field surveys of SCTLD, likely due to their small size, cryptic appearance, and fast progression of disease (making divers less likely to record active lesions). This study demonstrates that, like their larger adult counterparts, recruits are vulnerable to infection and can succumb relatively quickly (days to a few weeks), reducing the likelihood that their infection would be captured by surveys. In addition, protocols commonly used to identify disease on Caribbean reefs are not designed to examine juvenile corals. For instance, the Florida Reef Resilience Program’s Disturbance Response Monitoring survey, intended to “collect detailed monitoring data on diseased corals” (FWC, 2021), calls for a simple tally of corals ≤4 cm in diameter without space to document their condition (disease, recent mortality, etc.). Divers are also instructed not to count corals <1 cm in diameter, therefore missing many recruits <1 yr in age. Similarly, the Atlantic and Gulf Rapid Reef Assessment coral monitoring protocol asks divers to measure and report the status of colonies ≥4 cm in diameter (AGRRA, 2021). Without accounting for the smallest colonies, we have likely underestimated the true extent of SCTLD infection and mortality on Caribbean reefs.

Lesion appearance and progression in *C. natans* and *D. labyrinthiformis* recruits largely matched similar descriptions for adult colonies (FKNMS, 2018; Landsberg et al., 2020; Eaton et al., 2021). Beginning in some recruits of both species after just 48 h of exposure, lesions continued to present in new, previously healthy recruits over time. Lesions typically began at the periphery of a recruit before advancing across it, characterized by both focal and multifocal tissue loss that left behind exposed skeleton.

As adults, *C. natans* and *D. labyrinthiformis* are among the species that are most susceptible to SCTLD (Precht et al., 2016; Lunz et al., 2017; Alvarez-Filip et al., 2019), exhibiting high disease prevalence and rapid tissue loss (Meyer et al., 2019; Estrada-Saldívar et al., 2020; Meiling et al., 2021; Williams et al 2021). We observed high rates of tissue loss in *C. natans* recruits throughout exposure (26.3 ± 13.6% per day), with many dying within just two days of developing initial lesions, surpassing the average rate of ~10% reported for conspecific adults (Meiling et al. 2021). In contrast, lesions progressed much slower on average in infected *D. labyrinthiformis* throughout Dose 1 (2.9 ± 3.7% per day), similar to the rates that Meiling et al. (2021) reported for adult colonies of several species. However, it may not be accurate to directly compare rates of tissue loss between large, old, established adult colonies and small, single- or few-polyp recruits. Adults have thicker tissues and orders of magnitude more polyps than recruits, presumably giving them far greater energy reserves and resources to draw upon for defense (Barnes & Lough, 1992; Loya et al., 2001).

Five small *C. natans* recruits in the control treatment died during the experiment without exhibiting sloughing tissue or denuded skeleton, likely reflecting the generally low survivorship that characterizes the early life stages of corals in general (Vermeij & Sandin, 2008). None of these recruits showed signs of disease and instead appeared to be negatively affected by turf algae that grew around them on the ceramic plugs. In this case, their small size seemed to be a disadvantage, as there was no mortality recorded in larger recruits in the control treatment.

In addition to elucidating how Caribbean coral recruits present with SCTLD, this study allowed us to investigate relative versus absolute disease resistance. While no recruits managed to withstand infection during a second (presumably potent) dose of SCTLD, many were found to avoid infection during the first dose. After 20 days of SCTLD exposure, 59.0% of *C. natans* recruits had died compared with only 3.1% of *D. labyrinthiformis* recruits. Given our experimental design, we cannot distinguish whether the variable patterns of disease progression and mortality between the two cohorts of recruits were caused by differences in age, size, species, or a combination of these factors. Size may have contributed to this disparity, as the largest *C. natans* recruits were comparable in area to the smallest *D. labyrinthiformis* recruits (~10 mm^2^), and survivorship in *C. natans* recruits was significantly positively correlated with recruit area. Size has been shown to influence recruit survival in the face of other stressors, including ocean acidification, predation, and competition (Doropoulos et al 2012). However, Williams et al. (2021) found that SCTLD disproportionately affected the largest adult colonies in the field. This suggests that despite the benefits of larger size for recruits, there may come a point later in ontogeny when size becomes a disadvantage for corals exposed to SCTLD, perhaps because greater surface area results in higher chances of infection by waterborne pathogens as they encounter coral surfaces. Since coral immunity matures throughout early ontogeny (Frank et al 1997; Nozawa and Loya 2005), the four-month age difference between the *C. natans* and *D. labyrinthiformis* cohorts may also have contributed to relatively lower susceptibility of the older *D. labyrinthiformi*s. To inform restoration efforts using sexually-produced recruits, future studies should investigate whether SCTLD vulnerability varies as recruits age and grow. Such information may help managers minimize mortality downstream by optimizing the duration of recruit grow-out in protected nurseries or *ex-situ* facilities prior to outplanting on reefs in the disease-endemic zone (Randall et al., 2020).

Our results also provide evidence for the relatively higher resistance of chimeric colonies to SCTLD, at least for *C. natans*. This appears to be true both from the perspective of individual recruits (i.e., recruits benefit on an individual basis when grouped together) and from the perspective of the larger chimeric grouped entity (i.e., the emergent colony that results from the fusion of multiple individual recruits). In *C. natans*, being part of a group increased the probability of survival by up to 50% compared with single recruits of the same size (Fig. 7A). In addition, small and medium-sized chimeras were significantly more likely to survive SCTLD exposure (i.e., at least some part of the chimeric colony remaining apparently healthy) compared with single recruits of a similar size as the chimera (Fig. 8). In some cases, one member of a chimera would become infected while others remained apparently healthy, suggesting that being part of a group is not universally protective. However, rarely did all members of a chimera die following the infection of one member, particularly as the number of genets forming the chimera increased (Fig. 7B).

It has long been observed that coral larvae behave gregariously, resulting in aggregations of settlers that contain multiple genets and that are often spawned from different parents (Duerden, 1902; Frank et al., 1997; Hidaka et al., 1997; Raymundo & Maypa, 2004; Puill-Stephan et al., 2011; Puill-Stephan et al., 2012). The formation and fusion of multi-polyp chimeras during early ontogeny confer numerous benefits that seemingly outweigh the potential costs of conflict among partners (Amar et al., 2008). Chimeras have been shown to exhibit significantly higher survival and growth than individual recruits, and the immediate increase in size achieved through fusion helps vulnerable coral recruits rapidly surpass size-escape thresholds (Raymundo & Maypa, 2004; Amar et al., 2008; Christiansen et al., 2008; Vermeij & Sandin, 2008; Puill-Stephan et al., 2011; Doropoulos et al., 2016; Sampayo et al., 2020; Huffmyer et al., 2021). In addition, chimerism may be a powerful mechanism of “evolutionary rescue”, because the genetic variation present in multi-partner aggregations may enhance plasticity and stress tolerance compared with single-genet individuals (Rinkevich & Yankelevich, 2004; Rinkevich et al., 2016; Rinkevich, 2019; Huffmyer et al., 2021). Huffmyer et al. (2021) reported that fused *Pocillopora acuta* recruits survived five days longer than individuals under high temperature (+2.5 °C), and even longer when multiple parental genotypes were represented. In this study, the relative resistance to SCTLD observed in chimeric *C. natans* entities may be derived in part from their ability to draw upon a wider repertoire of resources and traits, including those involved in resisting disease. If this is the case, restoration efforts may target the formation of chimeras as a strategy to maximize recruit resilience in the face of SCTLD.

Despite the advantages of size and grouping observed in *C. natans*, these factors did not explain patterns of susceptibility in *D. labyrinthiformis*. Over half of the *D. labyrinthiformis* recruits exposed to SCTLD remained visually healthy during Dose 1, and the majority that became infected showed a stall in lesion progression after an initial wave of tissue loss (Fig. 4). All but one recruit survived Dose 1 (Fig. 4). These results are consistent with observations of apparent resistance in the field, where some colonies show halted tissue loss (or escape infection altogether) despite high SCTLD prevalence in adjacent colonies (Sharp et al. 2020). Such patterns suggest the existence of differentially resilient colonies or communities that fail to succumb despite tissue loss (and in some cases, death) in many of their neighbors (Sharp et al. 2020). Given that all *D. labyrinthiformis* recruits were reared under identical conditions, spawned from the same cohort of eight parents, and hosted *D. trenchii*, fine-scale genotypic variation may be responsible for their different disease responses.

The addition of a second, possibly more potent, dose of active SCTLD (Dose 2) revealed that no recruits of either cohort seemed to possess “absolute” resistance to SCTLD. This second treatment might have parallels in nature, representing an ongoing outbreak in which the contagious disease initially infects a few colonies with a relatively low dose but then spreads among colonies over time, progressively infecting more of the population (Williams et al., 2021) and resulting in a higher dose to any surviving colonies. In general, SCTLD appears to be much more prevalent than other coral diseases (Alvarez-Filip et al., 2019), with many colonies on a reef typically infected at a given time. Williams et al. (2021) reported that corals within 3 m of a diseased colony are at higher risk of contracting SCTLD than those further away. These findings suggest a plausible scenario in which recruits on a reef are located in close proximity to adult colonies that become infected one after the other, repeatedly exposing the recruits to pathogen loads that they may not be able to withstand.

Other SCTLD-susceptible species, such as *Orbicella faveolata* and *Montastraea cavernosa*, are major reef-builders throughout Florida and the Caribbean and are the target of reef restoration efforts using sexually-produced recruits. As such, juveniles of these species should be tested for rates of infection and tissue loss to assess the risk of SCTLD when outplanted in areas where it is endemic or emergent (Aeby et al., 2021). Since adult colonies of different species exhibit variation in their immunocompetency (Palmer et al., 2011) and rates of lesion progression (Meiling et al., 2021), they may also show varied responses to SCTLD as juveniles. In addition, understanding how SCTLD interacts with other stressors, such as elevated temperature, sedimentation, and nutrient pollution, may be important because they can increase disease susceptibility (Rodriguez-Lanetty et al., 2009; Vega Thurber et al., 2013; Pollock et al., 2014; Rädecker et al., 2015; Maynard et al., 2015; Zaneveld et al., 2016; Ward et al., 2017; Walton et al., 2018; Aeby et al., 2020; Howells et al., 2020).

Despite increasing attention to SCTLD over the past several years, relatively little effort has been made to create standardized protocols to test susceptibility (i.e., methods for exposing corals, how to establish or titrate a standardized dose, etc.). It is likely that the doses delivered in many laboratory experiments, including this one, may be higher (perhaps by orders of magnitude) than those typically experienced by corals in the field, due to the spread and dilution of a waterborne pathogen. In addition, many studies have used contact assays, which may be less accurate in reflecting mechanisms of SCTLD transmission on the reef. As such, researchers should collaborate to identify appropriate, ecologically relevant doses and delivery techniques to make SCTLD studies as replicable and relevant as possible.

Our results indicate that recruits are indeed susceptible to SCTLD and suggest that managers will need to mitigate the risk of disease in restoration efforts using coral recruits. Antibiotic treatment has been shown to be highly effective in halting lesion progression in adult colonies (Aeby et al., 2019; National Academies, 2019; Neely et al., 2020; Walker et al., 2021; Neely et al., 2021b; Shilling et al., 2021), but considering the rapid rates of tissue loss and mortality reported here, it may be extremely challenging for managers to detect lesions and apply antibiotics in time to save an actively diseased juvenile. Therefore, instead of combating disease once it appears, managers should consider preemptive approaches to prevent disease from infecting recruits. This could potentially be achieved through a variety of interventions, including: (1) holobiome manipulations of juveniles with the goal of decreasing SCTLD susceptibility, and/or (2) selective breeding of new generations that are more resistant to SCTLD using parents that are more resistant.

Holobiome manipulations include modifying the communities of algal endosymbionts (Family Symbiodiniaceae) in coral tissue (van Oppen et al., 2015). Landsberg et al. (2020) reported that SCTLD involves a disrupted association between a host coral and its endosymbionts, suggesting that susceptibility may be attributed to differences between algal genera in addition to coral species. Dennison et al. (2021) found that corals hosting *Breviolum*, both as their dominant symbionts and at background levels, were more likely to become infected with SCTLD than corals hosting other genera. In addition, inshore corals hosting *Durusdinium trenchii* have largely escaped the impacts of SCTLD relative to their offshore counterparts hosting members of the genus *Breviolum* (Rubin et al. 2021). However, since almost all recruits in this study hosted primarily *D. trenchii*, we were not able to assess the influence of Symbiodiniaceae identity on disease incidence. Future studies should test the role of different endosymbiont taxa in recruit susceptibility. If a specific taxon (i.e. *Breviolum*) is implicated in SCTLD infection, managers may choose to provide recruits with sources of alternative taxa during initial symbiosis establishment (Williamson et al., 2021).

Within the diverse array of prokaryotes that constitute a coral’s microbiome, some members have been shown to promote holobiont health and aid in the removal of sources of stress (i.e. pathogens, reactive oxygen species) (Nissimov et al., 2009; Shnit-Orland & Kushmaro, 2009; Krediet et al., 2013; Glasl et al., 2016; Peixoto et al., 2021). For example, Raina et al. (2016) isolated large quantities of the antimicrobial compound tropodithietic acid (TDA) produced by coral-associated *Pseudovibrio* bacteria, which prevented the growth of pathogens. Active delivery and/or facilitation of such beneficial microorganisms for corals (BMCs) in the form of probiotics have the potential to minimize impacts of various stressors on adult corals (Peixoto et al., 2017; Rosado et al., 2019; Peixoto et al., 2021). In a controlled laboratory experiment, inoculations with a consortium of BMCs partially mitigated bleaching and pathogen exposure in *Pocillopora damicornis* (Rosado et al., 2019). Given the apparent success of these treatments in adult corals, managers should consider dosing juveniles with probiotics or BMCs as an adaptive intervention prior to outplanting to reduce their risk of contracting SCTLD. However, associations with both prokaryotic and eukaryotic microbes can be transient, shifting with ontogeny and environmental conditions and thus offering only short-term protection against disease (Little et al., 2004; Abrego et al., 2009; Sharp et al., 2012; Hernández-Agreda et al., 2016; Damjanovic et al., 2017; Cumbo et al., 2018).

In order to promote longer-term resistance in new generations of corals, future studies should investigate the heritability of SCTLD susceptibility and resistance. The recruits in this study came from parents that were “rescued” from the Dry Tortugas ahead of the disease front as part of the Florida Coral Rescue. As such, they had never been exposed to SCTLD, and can be considered naive to the disease and therefore likely susceptible. In contrast, colonies that have survived in the endemic zone for several years may harbor some degree of resistance, and managers should consider utilizing them for both asexual propagation and managed breeding (assisted gene flow) to ensure the persistence of susceptible species (Baums et al., 2019). Since other traits such as heat tolerance can be passed on to the next generation (Quigley et al., 2020; Howells et al., 2021), perhaps there are heritable components of disease resistance in corals. Selectively breeding colonies that have persisted in the endemic zone may increase the frequency of SCTLD-resistant alleles in offspring, improving outcomes in the face of disease while enhancing genetic diversity in dwindling populations (National Academies, 2019; Baums et al., 2019; Voolstra et al., 2021).

Since its initial outbreak in Florida, SCTLD has spread rapidly throughout the Caribbean and caused extensive mortality among susceptible species (Alvarez-Filip et al., 2019; Kramer et al., 2019; Estrada-Saldívar et al., 2020; Brandt et al., 2021; Costa et al., 2021; FDEP, 2021a; Dahlgren et al., 2021; Heres et al., 2021). In light of the considerable infection and mortality of recruits reported here, we can infer that untold numbers of coral recruits have likely been lost as a result of SCTLD outbreaks. Although these casualties have gone undocumented, they nevertheless impact the potential for species and reef recovery. Over the past several decades, the number of juvenile corals on Caribbean reefs has already declined significantly (Vermeij et al., 2011). If the few recruits that remain cannot survive and grow because they succumb to disease, they cannot contribute to ecosystem functioning and the restoration of damaged reefs (Miller et al., 2000; Miller & Barimo, 2001; van Woesik et al., 2014).

Entire regions have now become affected by SCTLD, with outbreaks spanning up to 140 km in scale (Muller et al., 2020). Such a large “footprint of disturbance” threatens the persistence of susceptible species, with larvae from unaffected areas needing to disperse over greater and greater distances to reach and replenish depleted reefs (Dietzel et al., 2021). The recovery of dwindling populations will require both the survival of mature colonies as broodstock and the recruitment of juveniles (Hughes & Tanner, 2000). If SCTLD outbreaks continue to reduce the number of spawning adults and infect recruited offspring early on, the potential for natural population recovery may be severely limited. For this reason, multiple, active intervention strategies may be critical to avoid extirpation of susceptible species and ecosystem decline on Caribbean reefs.

## Acknowledgements

The authors thank The Florida Aquarium’s Center for Conservation for providing the coral larvae that were later utilized in this experiment. We also thank the Florida Fish and Wildlife Commission for providing the parent colonies that spawned at The Florida Aquarium through the Coral Rescue Project. Parent corals were collected under FKNMS Superintendent’s permit FKNMS-2017-100, and larvae were transferred to the University of Miami under permit an Animal Transaction Agreement with the Florida Aquarium (USDA LIC# 58-C-0977, AZA# 2305000).

